# MassiveFold data for CASP16-CAPRI: a systematic massive sampling experiment

**DOI:** 10.1101/2025.05.26.653955

**Authors:** Nessim Raouraoua, Marc F. Lensink, Guillaume Brysbaert

## Abstract

Massive sampling with AlphaFold2 has become a widely used approach in protein structure prediction. Here we present the MassiveFold CASP16-CAPRI dataset, a systematic, large-scale sampling of both monomeric and multimeric protein targets. By exploiting maximal parallelization, we produced up to 8,040 models per target and shared them with the community for collaborative selection and scoring. This collective effort minimizes redundant computation and environmental impact, while granting resource-limited groups - especially those focused on scoring - access to high quality structures. In our analysis, we define an interface-difficulty classification based on DockQ metrics, showing that massive sampling yields the greatest gains on most of the challenging interfaces. Crucially, this classification can be predicted from the median ipTM scores of a routine AF2 run, enabling users to selectively deploy massive sampling only when it is most needed. Combined with a reduction of the massive sampling from 8,040 to 2,475 predictions, such targeted strategies dramatically cut computation time and resource use with minimal loss of accuracy. Finally, we underscore the persistent challenge of choosing optimal models from massive sampling datasets, emphasising the need for more robust scoring methods. The MassiveFold datasets, together with AlphaFold ranking scores and CASP and CAPRI assessment metrics, are publicly available at https://github.com/GBLille/CASP16-CAPRI_MassiveFold_Data to accelerate further progress in protein structure prediction and assembly modeling.

## Introduction

Since its debut with AFsample^1^, massive sampling with AlphaFold2^2,3^ has consistently outperformed alternative approaches, especially for the prediction of protein complexes. In CASP15-CAPRI, the Wallner group’s top ranking demonstrated that pushing AF2 for greater sampling and diversity can uncover solutions otherwise missed^4–6^. We then refined this strategy in MassiveFold^7^, leveraging maximal parallelization on GPU clusters to reduce turnaround times.

Although MassiveFold is very efficient and easy-to-use, its biggest hurdle remains selecting the best models from its vast output^8^. Relying almost exclusively on AF2’s internal scores, mainly ipTM for multimers, limits performance, especially on targets where AF2’s confidence estimates are unreliable. Ideal scoring strategies would be purely structure-based and independent of AF2 itself; this would empower the wider structure-prediction community to exploit the full potential of deep-sampling strategies.

To that end, we transformed model selection into a collaborative effort by rendering MassiveFold-produced models available to participants in CASP16’s stage 2, QMODE3 and CAPRI’s scoring challenge. For the standard stage 1, we submitted our top five models - chosen by AF2 confidence - from the up to 8,040 MassiveFold predictions, under the ‘Brysbaert’ group. We then made the entire dataset available to all participants for stage 2, allowing them to select from a potentially enriched set of structures. Our goal was to create a single, shared massive sampling dataset so that groups who simply wanted to run AF2 extended sampling (possibly with the dropout-based diversity options as introduced by AFsample) can skip the heavy lifting themselves, in response to a pattern noted in previous CASP and CAPRI experiments. Although CASP stage 1 still encourages groups to perform their own extended runs, pooling massive sampling efforts enhances collective engagement and empowers both scoring experts and routine AF2 users to tap into a rich, pre-computed library of models.

We generated MassiveFold data for both monomers and multimers. While monomer prediction is largely solved by AF2, the question remains if massive sampling can probe alternative conformations. Our primary goal, however, was to spur development of better scoring algorithms for protein interfaces in multimeric assemblies. Accordingly, this manuscript focuses on those multimeric targets, specifically the ones offered for blind prediction by both CASP and CAPRI. However, all MassiveFold data with AlphaFold2 ranking scores and assessment metrics are publicly available for any downstream use.

## Materials & Methods

### Targets

We predicted the structures of CASP16 & CAPRI Round 57 targets. The full list of targets can be found on the CASP16 target page of the Prediction Center (https://predictioncenter.org/casp16/targetlist.cgi?view_targets=all). We selected only those targets consisting of protein chains, resulting in 32 monomeric and 39 multimeric targets, for a total of 71 targets. For the purpose of this manuscript, we focused on a subgroup of targets, namely the multimeric targets that were shared between CASP16 and CAPRI Round 57. These 31 common targets were then subdivided into 65 unique interfaces, which were individually evaluated by the CAPRI assessors. We focused on these interfaces for our analysis.

### Production of the massive sampling sets

Structure predictions were generated using MassiveFold v1.2.3 on the Jean Zay supercomputing cluster at IDRIS (Institut du Développement et des Ressources en Informatique Scientifique) in Paris, France, employing V100 and A100 GPUs. For most targets, a standard set of 8,040 distinct structure predictions was generated. However, for a selected few assemblies this number was reduced due to the target size demanding prohibitively high computational efforts (6/39 for CASP, 2/31 for CAPRI).

The prediction volume was obtained by systematically combining several sources of diversity: inference tools, neural network (NN) models, and unique parameter sets. These parameter sets were initially inspired by the Wallner parameterization used in CASP15^4^ and subsequently refined during our participation in CAPRI Rounds 55 and 56^8^. At the time of CASP16, MassiveFold incorporated two primary inference tools: AFmassive^7^ (a modified version of AlphaFold2 optimized for massive sampling) and ColabFold^9^ (leveraging the ColabFold_DB and MMseqs2 alignment approach for added diversity). Each parameter set was devised to produce 1,005 structures by systematically sampling across the available NN models. The specific sampling strategy differed between multimers and monomers due to differences in availability of NN models. For multimers, we used 15 distinct NN models - 3 variant versions each comprising 5 base models^3^. By sampling 67 structures per model, we obtained the desired 1,005 predictions per parameter set (15×67). For monomers, only 5 NN models are available, as DeepMind released only a single version for monomeric targets. To match the same output size, we sampled 201 structures per model (67×3), again yielding 1,005 predictions per set. In total, the standard set of 8,040 predictions per target was assembled from 6 parameter sets run using AFmassive and 2 sets run with ColabFold. The specific conditions for each set are summarized in Table 1.

**Table 1:** Parameter sets used for massive sampling with MassiveFold in CASP16-CAPRI.

| Setup | Dropout<br>Evoformer | Dropout structure<br>module | Templates | Recycles | Structure inference<br>engine |
| --- | --- | --- | --- | --- | --- |
| afm_basic |  |  | X | 21 | AFmassive |
| afm_woTemplates |  |  |  | 21 | AFmassive |
| afm_dropout_full | X | X | X | 21 | AFmassive |
| afm_dropout_full_woTemplates | X | X |  | 21 | AFmassive |
| afm_dropout_full_woTemplates_r3 | X | X |  | 3 | AFmassive |
| afm_dropout_noSM_woTemplates | X |  |  | 21 | AFmassive |
| cf_woTemplates |  |  |  | 21 | ColabFold |
| cf_dropout_full_woTemplates | X | X |  | 21 | ColabFold |

The produced data was used in three ways. First, when a target and its stoichiometry were released for stage 1 predictions, we ran MassiveFold to generate 8,040 structure predictions. From this set, we selected the top five models for our ‘Brysbaert’ group submission, based on AlphaFold confidence scores; this selection can thus be considered a massive sampling baseline (see next section). Second, the full dataset was shared with the CASP-CAPRI community via the PLBS/SINBIOS platform (https://sinbios.plbs.fr/) for data storage, enabling its use in CASP stage 2, QMODE3, and the CAPRI scoring phase. The data remained hosted there throughout the experiment. Third, following the conclusion of CASP16, we moved the datasets to long-term storage, assigning each target a DOI within the CASP16-CAPRI MassiveFold collection hosted on the “Recherche Data Gouv” repository (https://entrepot.recherche.data.gouv.fr/dataverse/casp16mf). Additionally, the 25 default predictions from the AlphaFold2-v3 run were extracted from the MassiveFold set, defining the AF2-baseline set.

### Prediction selection and assessment

In order to select the predictions for submission, we used the ranking confidence, which is common to AlphaFold2 and ColabFold. For monomers, this is computed as the mean pLDDT calculated over the individual residue pLDDT values; it ranges from 0 to 100 and can therefore be considered a confidence percentage. For multimers, this is computed as 0.8*ipTM+0.2*pTM, where pTM is a predicted TM-score and ipTM is the pTM computed between chains. These scores range from 0 to 1. In both cases a higher score indicates more confidence. We submitted the five highest-confidence predictions out of the whole massive sampling set as the ‘Brysbaert’ predictor submission. CASP assessment of the MassiveFold data was performed in the QMODE3 stage; the data are available on the CASP website (https://predictioncenter.org/). Using the native structure, we also calculated the CAPRI metrics^6,10^ for all models and used these in our subsequent analyses.

### Assignment of interface difficulty

We define three levels of interface difficulty: ‘easy’, ‘hard’ and ‘extreme’. These reflect the ease with which MassiveFold could generate predictions of acceptable quality or better, following CAPRI definitions. This classification considers the full DockQ score distribution, using two key metrics: the maximum DockQ (DockQ_max_) and the spread of the top distribution, DockQ_max-Q3_ computed as DockQ_max_ minus the third quartile. The thresholds used to classify the 65 interfaces are listed here and shown in Figure **1ab**:

**Figure 1:**
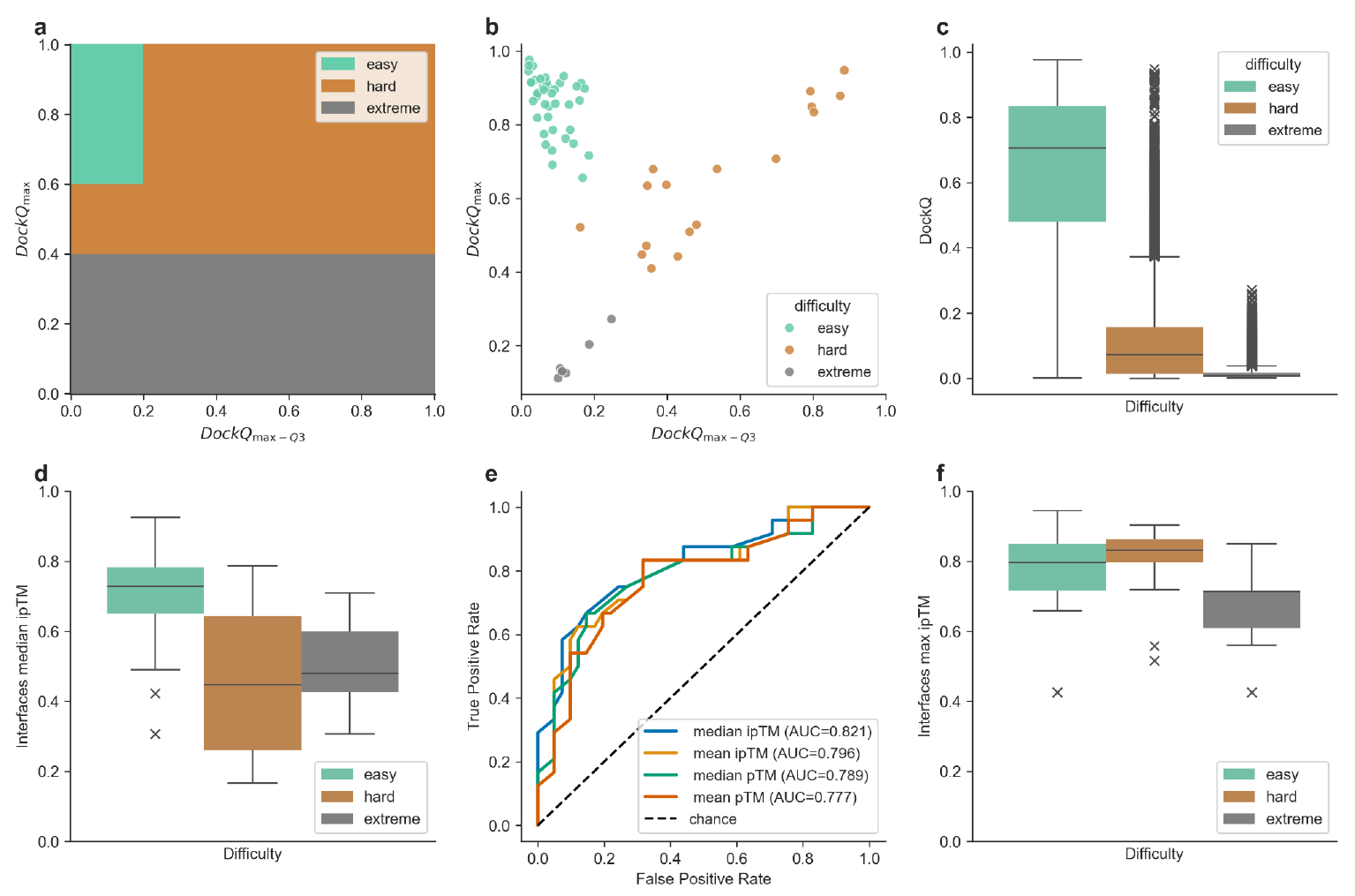
Interfaces assigned difficulty levels, applying the following color code: green for ‘easy’, brown for ‘hard’ and gray for ‘extreme’. **a**) illustrates the thresholds of DockQ_max_ and DockQ_max-Q3_, used to determine these difficulty levels following the interfaces DockQ-based metrics distribution of **b**). **c**) boxplots of all predictions’ DockQ values, per difficulty category. **d**) boxplots of the AF2-baseline median ipTM for each interface and per difficulty class (25 AF2-v3 predictions per target). **e**) Receiver operating characteristic (ROC) analysis for classification into ‘non-trivial’ targets (‘hard’ and ‘extreme’) using different AF2-based metrics with their respective area under the curve (AUC) values. **f**) boxplots of the max ipTM of each interface in the final massive sampling set per difficulty. All boxplots extend from the first quartile to the third quartile, with a line at the median; the whiskers reach out to the furthest data point still within 1.5 times the interquartile range from the box, and outliers beyond the whiskers are shown as crosses.

- ‘Easy’ interfaces (n=41): (DockQ_max_ > 0.6) & ((DockQ_max-Q3_) ≤ 0.2)
- ‘Hard’ interfaces (n=17): (0.4 < DockQ_max_ < 0.6) or ((0.6 < DockQ_max_ < 1) & ((DockQ_max-Q3_) ≥ 0.2))
- ‘Extreme’ interfaces (n=7): DockQ_max_ < 0.4

### Massive sampling scenarios

To selectively apply massive sampling to challenging interfaces, we grouped those labeled ‘hard’ and ‘extreme’ under the ‘non-trivial’ category. Since interface difficulty was defined based on the DockQ distribution of predictions, it could only be assessed after generating and evaluating the interfaces’ full MassiveFold dataset. To enable prediction of difficulty *a priori*, we analyzed the AF2-baseline confidence metrics, identifying the median ipTM score as the best predictor (AUC=0.821; Fig. **1e**). Based on this, we defined three sampling strategies - ranging from most to least computing-intensive - using different median ipTM thresholds (Fig. **1d**) to decide whether additional massive sampling should be performed. The first strategy - MassiveFold-S1 - computes massive sampling for any interface with AF2-baseline median ipTM below 0.8, excluding only a few ‘easy’ interfaces from massive sampling. MassiveFold-S2 excludes most ‘easy’ interfaces but also some ‘hard’ interfaces, by setting the threshold at 0.65, the value that separates the core distribution of these two categories. MassiveFold-S3, finally, uses 0.57 as a threshold which optimizes the F1-score for the median ipTM that has the best AUC (Fig. **1e**). We abbreviate these strategies to MF-S1, MF-S2 and MF-S3.

Aside from the difficulty prediction of interfaces *a priori*, we explored another method of sampling reduction. This method is based on the sampling analysis (Fig. **S1**) both in terms of intensity and diversity. We realised that the major benefits for ‘hard’ targets can be found in the first half of the sampling (Fig. **S1a**), with only marginal improvements in the second half, the interface of T296.1 being the only exception. We thus considered sampling 33 predictions per neural network model instead of the initial 67, which would decrease the total number of predictions per target from 8,040 (8 parameter sets x 15 NN models x 67 predictions) to 3,960 (8×15×33). Moreover, the evaluation of each individual run performance (Fig. **S1cdef**) reveals that ‘afm_dropout_full_woTemplates_r3’, the only run with 3 recycling steps (vs. 21 in all others), is extremely inefficient both on ‘easy’ and ‘hard’ interfaces and that ‘cf_dropout_full_woTemplates’ gets comparably poor results on ‘hard’ interfaces, with a low amount of ‘high’ quality predictions (Fig. **S1d**). To refine our choice of parameter sets, we identified ‘afm_dropout_full_woTemplates’ and ‘afm_dropout_noSM_woTemplates’ runs as having highly similar results, confirming previous results^8^, except on interface T296.1 where the latter performed uniquely well (DockQ ≈ 0.9 vs. other runs <0.6). For these reasons, we kept the 5 runs judged as essential by removing the 3 least performant or diverse ones being ‘afm_dropout_full_woTemplates_r3’, ‘cf_dropout_full_woTemplates’ and ‘afm_dropout_full_woTemplates’.

Combining the reduction of sampling intensity and diversity sources, the massive sampling size goes from an initial 8,040 (8×15×67) to 2,475 (5×15×33) predictions per ‘hard’ target. By using this method without or alongside the previous ipTM median threshold, we create 3 new scenarios: S4, S5 and S6. For S4, no interfaces are excluded from massive sampling, S5 uses the same threshold as S2, which is ipTM median < 0.65 and S6 uses the same as S3, median ipTM < 0.57.

### Data analysis

Predictions with more than 300 clashes were excluded from our MassiveFold set, as well as predictions that are considered as ‘clash’ according to CAPRI calculation for every participant including our group ‘Brysbaert’. All analyses were performed with Python 3, using the matplotlib 3.9.3, seaborn 0.13.2, panda 2.2.2 and scikit-learn 1.6.1 packages.

### Code and data availability

All CASP16-CAPRI MassiveFold data, together with CASP and CAPRI assessment metrics and AlphaFold confidence ranking are accessible from the repository associated to this article: https://github.com/GBLille/CASP16-CAPRI_MassiveFold_Data.

## Results and discussion

### Difficulty classes

We assigned a difficulty level to each predicted interface based on the complete set of MassiveFold predictions. As shown in Figure **1ab**, this classification relies on two metrics: the quality of the best prediction (DockQ_max_), and its relative improvement over the rest of the predictions (DockQ_max-Q3_). The 41 interfaces labeled as ‘easy’ generally do not need massive sampling, as AF2 already produces good-to-excellent quality models. The ‘hard’ class (17 interfaces) is where massive sampling proves most beneficial, revealing improved models that are outliers in the prediction distribution (Fig. **1c**). The ‘extreme’ class (7 interfaces) includes the most difficult interfaces, for which even massive sampling rarely yields acceptable models - only one such case was successful.

The AF2-baseline median ipTM appears to be a reliable indicator of an interface’s difficulty (Fig. **1de**) and can help determine whether a target should be submitted to massive sampling. As ‘easy’ targets benefit only marginally from extensive sampling (see next section), using the AF2-baseline median ipTM as a pre-selection criterion offers a practical and efficient strategy (see Massive sampling scenarios in Materials and Methods and below for threshold suggestions). Additionally, the maximum ipTM value within a MassiveFold set provides useful information *post hoc*; a threshold of ipTM = 0.72 effectively distinguishes ‘extreme’ interfaces from the ‘hard’ ones, by separating the third quartile of ‘extreme’ target from the lowest of the ‘hard’ quartiles (Fig. **1f**). Therefore, getting maximum ipTM < 0.72 after massive sampling correlates with a high probability for it to be an ‘extreme’ target. However, this observation should be interpreted cautiously due to the limited number of ‘extreme’ cases.

### MassiveFold prediction quality

We used CAPRI metrics to rank predictors and scorers on their performance for each of the three classes of interface (Fig. **2** and **S2**). For the 17 ‘hard’ interfaces (Fig. **2**), simply selecting the top five models by AF2 confidence from the 8,040 MassiveFold predictions (the ‘Brysbaert’ approach) outperformed the best possible picks from the AF2-baseline set (‘AF2-baseline-best’). However, this advantage vanishes for the other difficulty levels (Fig. **S2**). For ‘easy’ interfaces, higher quality models come from the AF2-baseline set rather than from the highest AF ranking confidence in the massive sampling dataset and submitted by the ‘Brysbaert’ group (Fig. **S2**, top). This suggests that, for ‘easy’ interfaces, carefully scoring and selecting the best predictions from a standard AF2 approach is more effective than massive sampling (see also Fig. **S5d**). For ‘extreme’ interfaces, neither massive sampling nor standard AF2 methods produced consistently acceptable models (Fig. **S2**, bottom). These ‘blind spots’ affect most participants and suggest that additional strategies - enhanced sampling or the use of alternative methods (*e*.*g*., AlphaFold3) - are needed. Using the maximum ipTM on the massive sampling may identify those targets (Fig. **1e**).

**Figure 2:**
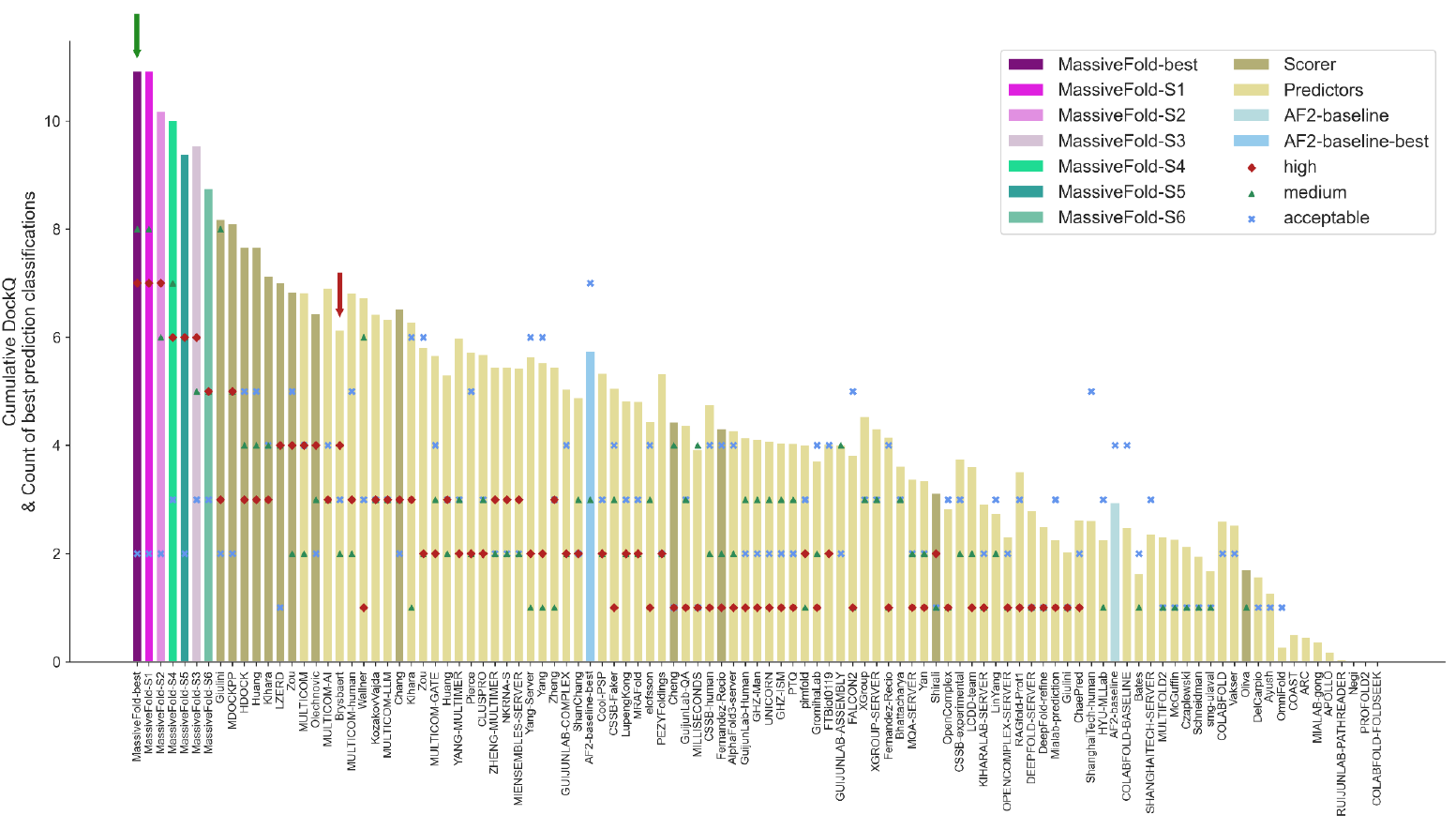
Ranking of groups that participated as predictors or scorers in CASP16-CAPRI for ‘hard’ interfaces. Each group’s ranking is based on their best prediction among the five submitted structures for each interface. The height of each bar represents the cumulative DockQ score across all interfaces, while the bars themselves are ordered according to the best CAPRI classification per interface, calculated as: 3×high+2×medium+acceptable. The counts for each classification are also indicated (high: red diamonds, medium: green triangles, acceptable: blue crosses). In addition to the official predictor and scorer participants, we included results from our massive sampling set: MassiveFold-best (the best DockQ structure from the entire set), and six simulated sampling strategies (MassiveFold-S1, S2, S3, S4, S5 and S6). For comparison, we also show AF2-baseline-best (the best prediction among the first 25 AF2v3 predictions per interface) and AF2-baseline (the prediction with the highest ipTM score).

Figure **2** shows that for ‘hard’ targets, the MassiveFold pool contains the largest number of high and medium-quality models (‘MassiveFold-best’). Yet relying solely on AlphaFold’s confidence ranking (as used by ‘Brysbaert’) fails to retrieve all of them (see also Fig. **S5c)**. This underscores the need for a dedicated scoring method to identify these buried gems; our MassiveFold dataset, complete with AlphaFold scores and CASP and CAPRI assessment metrics for every model, provides an ideal benchmark for developing such methods.

### Computing resources vs. performance trade-off

Classifying interfaces as ‘easy’, ‘hard’ and ‘extreme’ enabled us to evaluate the relevance of massive sampling for individual targets. Leveraging the AF2-baseline median ipTM (Fig. **1d**), we modeled three first gradual sampling strategies - scenarios MassiveFold-S1, S2, and S3 - which progressively reduce compute effort (Fig. **S4**, see Materials and Methods). These strategies are filtered subsets of ‘MassiveFold-best’, which incorporates all predictions. In MF-S1 (AF2-baseline median ipTM <= 0.8), we exclude only the easiest cases from massive sampling and see no drop in prediction quality while trimming runtime (Fig. **2, S3, S4 and S5a**). MF-S2 (AF2-baseline median ipTM <= 0.65) and MF-S3 (AF2-baseline median ipTM <= 0.57) further lighten the computational load, but due to the overlap in median ipTM values for ‘easy’ and ‘hard’ interfaces, these sampling scenarios occasionally misclassify a ‘hard’ interface as ‘easy,’ which leads to a reduced performance on those interfaces (Fig. **2**). Case in point: MF-S2 loses two ‘medium’ quality predictions, while MF-S3 loses one ‘high’ and three ‘medium’ quality predictions. Despite these minor losses, both strategies still rank above the other groups, while achieving substantial savings in GPU time (Fig. **2** and **S4**).

To go further in diminishing GPU computing time, we reduced the sampling intensity by half from 67 to 33 predictions per NN model per parameter set and the sampling diversity by keeping only 5 diversity parameter sets out of the 8 initially defined. This sampling trimming was combined with the filtering of the targets from MassiveFold-best, MassiveFold-S2 and MassiveFold-S3 to respectively establish the scenarios S4, S5 and S6 (Fig. **S4, S5b**, see Materials and Methods). These 3 scenarios still produce better quality models than any other participants (Fig. **2**). The cumulative DockQ between MF-S2 and MF-S4 are very close, however MF-S4 saves substantial computing time over MF-S2, and even over MF-S3 (Fig. **S3, S4**). The computing load is further decreased with scenarios MF-S5 and MF-S6 to more than five times from MF-best to MF-S6, although at the cost of certain ‘medium’ and ‘high’ quality models. MF-S5 (AF2-baseline median ipTM <= 0.65, 33 predictions per NN model and 5 diversity parameter sets) is a good compromise, retaining a high number of ‘medium’ and ‘high’ quality models compared to the other scenarios, and reducing the computing load about four times compared to MF-Best.

A comparison of cumulative DockQ distributions against the AF2-baseline (Fig. **S3**) underscores the urgent need for a robust scoring function that is able to identify the best models, even within standard AF2 runs (‘AF2-baseline-best’ compared to ‘AF2-baseline’).

## Conclusion

In this work, we generated and publicly released exhaustive massive-sampling datasets for both monomeric and multimeric targets in the CASP16-CAPRI experiment. Across 65 interfaces, massive sampling enabled top-rank predictions for all but seven ‘extreme’ cases. Its greatest benefit was observed on ‘hard’ interfaces, where it consistently produced one of the highest-quality models compared to the other participants. Crucially, we demonstrate that interface difficulty can be predicted *a priori* and with high accuracy from the median ipTM value of a standard AF2 run, allowing massive sampling to be deployed selectively and efficiently. This targeted approach combined with considerably less predictions in a massive sampling strategy achieves significant reductions in computation time and resource consumption, with only minimal impact on prediction quality. Moving forward, the principal challenge lies in devising a robust, preferably structure-only based scoring function, which allows to capitalize on the advantages of massive sampling in protein structure prediction.

## Acknowledgements

The French State under the France-2030 programme and the Initiative of Excellence of the University of Lille are acknowledged for the funding and support granted to the R-CDP-24-002-PIE project. This work was granted access to the HPC resources of IDRIS under the allocations 2024-AD010715332, 2024-AD010715333 and 2024-AD010715407 made by GENCI (Grand Equipement National de Calcul Intensif). We also acknowledge the University of Lille’s Intensive Scientific Computing Mesocentre, the Bilille platform and the SINBIOS platform for providing access to computing resources and storage space for the project. Finally, we thank Research Data Gouv and CNRS Research data for long-term hosting of the data.

