## Supplementary Material for "MassiveFold data for CASP16-CAPRI: a systematic massive sampling experiment"

### Supplementary Figures

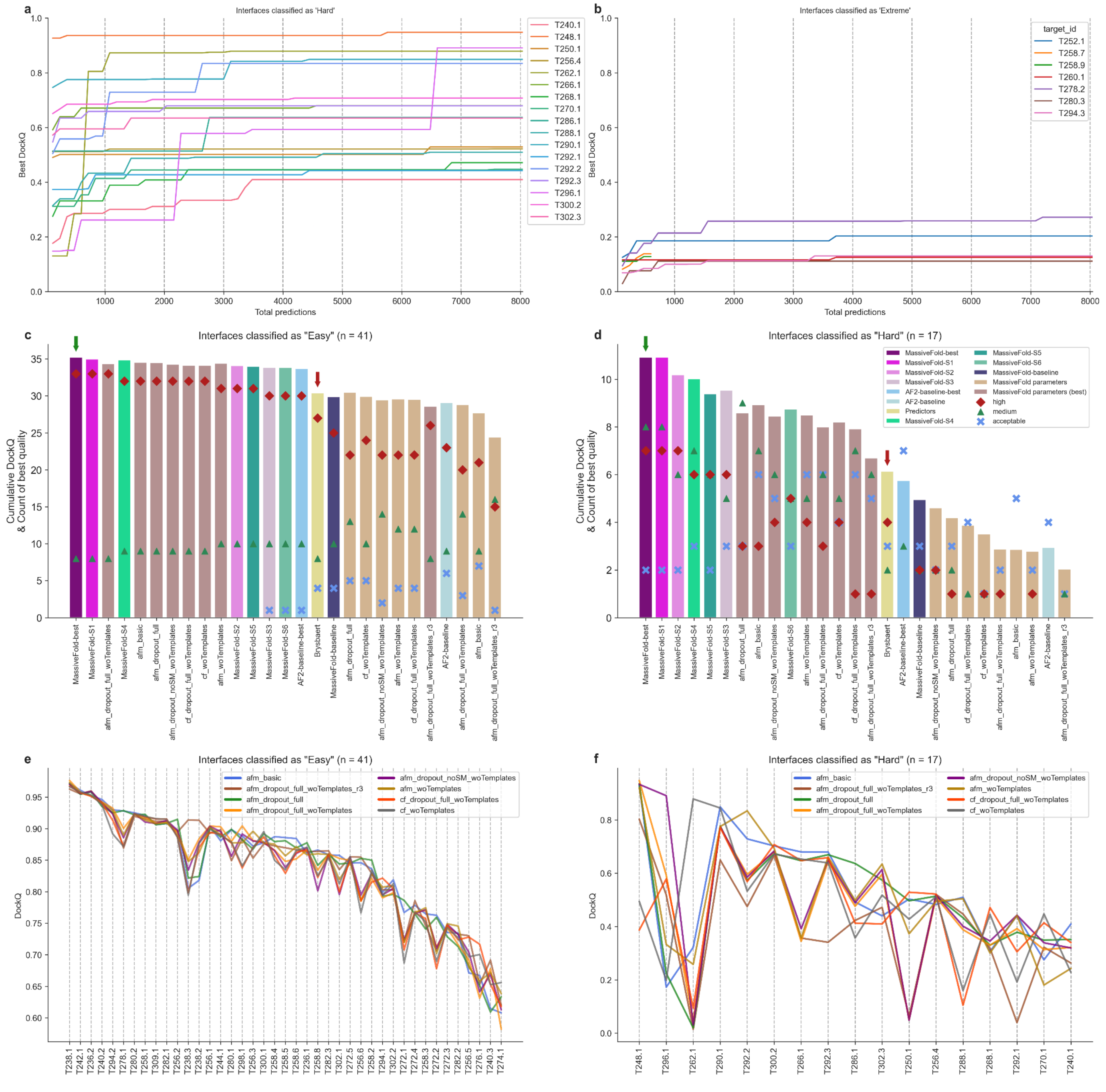


**Figure S1**: Analysis of both the intensity (**ab**) and the diversity (**cdef**) of the massive sampling. The best DockQ as a function of the number of predictions produced for 'hard' interfaces is illustrated in **a**) and 'extreme' ones in **b**). The parameter sets and the sampling scenarios are ranked for comparison in **c**) for 'easy' interfaces and **d**) for 'hard' ones. The interfaces best DockQ per parameter set is plotted for 'easy' interfaces in **e**) and 'hard' ones in **f**), interfaces are ordered by descending order of best DockQ.

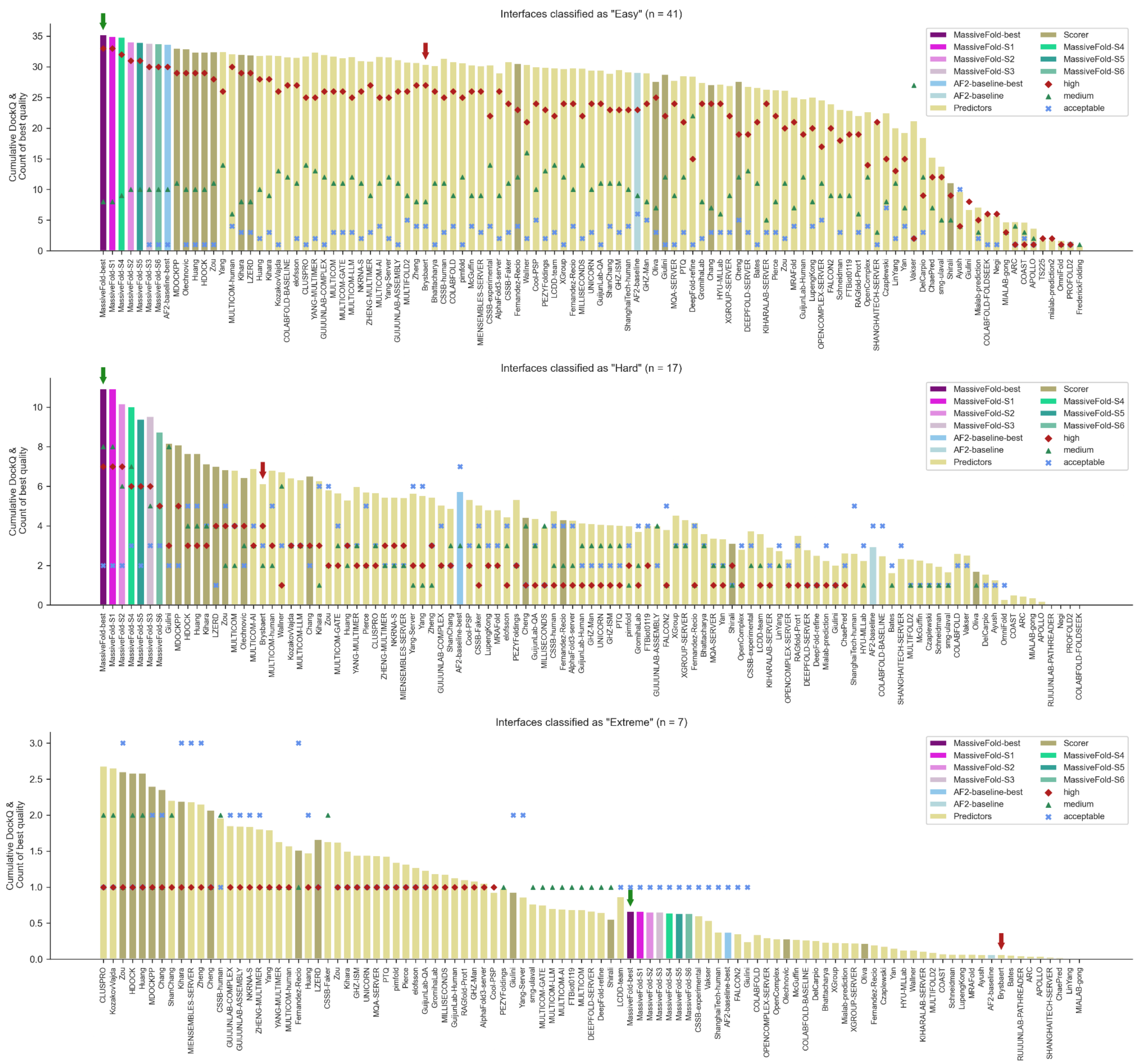


**Figure S2:** Ranking of groups that participated as predictors or scorers in CASP16-CAPRI for 'easy' (top), 'hard' (middle) and 'extreme' interfaces (bottom). Each group's ranking is based on their best prediction among the five submitted structures for each interface. The height of each bar represents the cumulative DockQ score across all interfaces, while the bars themselves are ordered according to the best CAPRI classification per interface, calculated as: 3×high+2×medium+acceptable. The counts for each classification are also indicated (high: red diamonds, medium: green triangles, acceptable: blue crosses). In addition to the official predictor and scorer participants, we included results from our massive sampling set: MassiveFold-best (the best DockQ structure from the entire set), and six simulated sampling strategies (MassiveFold-S1, S2, S3, S4, S5 and S6). For comparison, we also show AF2-baseline-best (the best prediction among the first 25 AF2v3 predictions per interface) and AF2-baseline (the prediction with the highest ipTM score among the first 25 AF2v3 predictions).

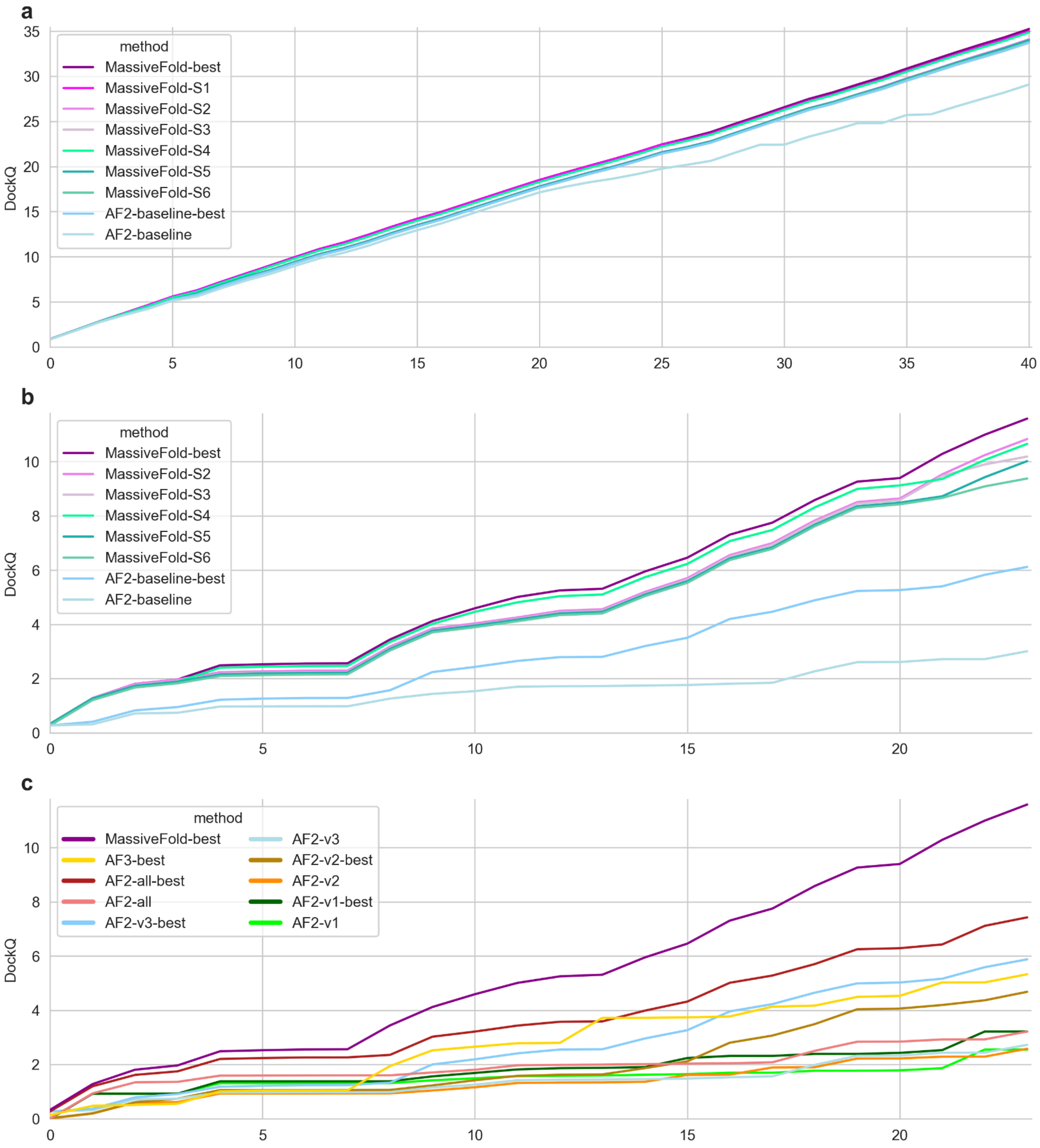


**Figure S3:** Cumulative DockQ depending, in **a**) and **b**), on the massive sampling strategy used, and in **c**), on the tool-selection combination. The interfaces are separated by difficulty with 'easy' in **a)** and non-trivial ('hard' and 'extreme') in **b**) and **c)**. For comparison, we also show AF2-baseline-best (the best prediction among the first 25 AF2v3 predictions) and AF2-baseline (the prediction with the highest ipTM score among the first 25 AF2v3 predictions). **c**) Comparison of the different AF2 versions, either with selecting the best DockQ (with '-best') or the best ipTM (without '-best'). 'MassiveFold-best' is put as a reference. AF3-best is the best DockQ of the 'AlphaFold3-server' predictor.

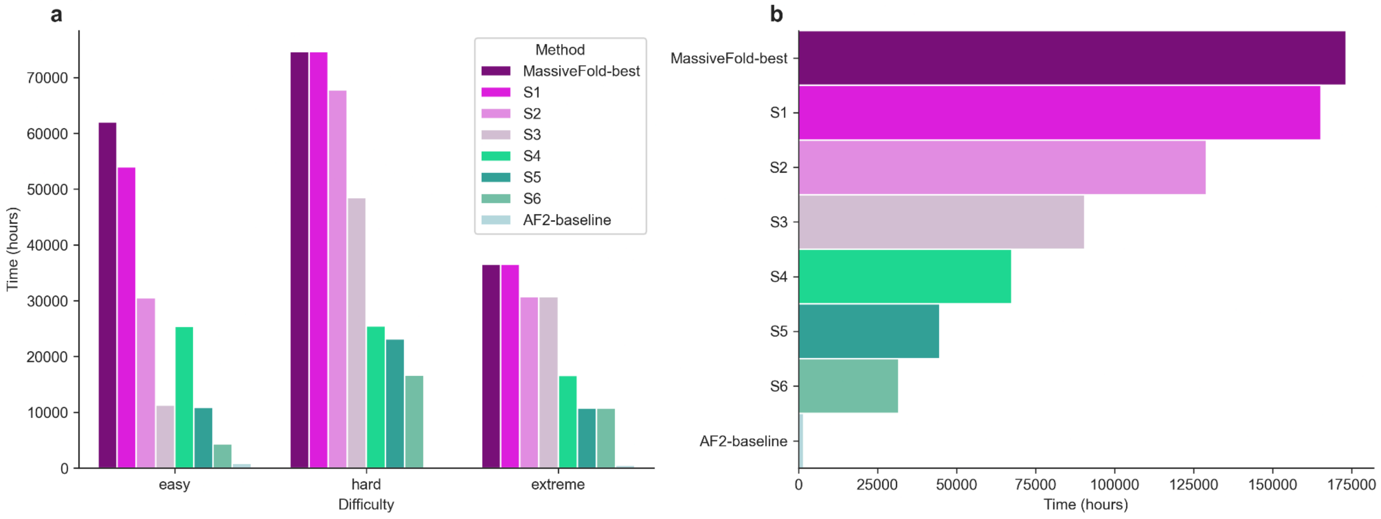


**Figure S4:** GPU time to compute MassiveFold data depending on the sampling strategy. **a)** Shows a comparison of these computing times per interface difficulty category between the different sampling strategies. **b)** Time to compute every interface. The AF2-baseline (25 first AF2v3 predictions) is also shown for comparison.

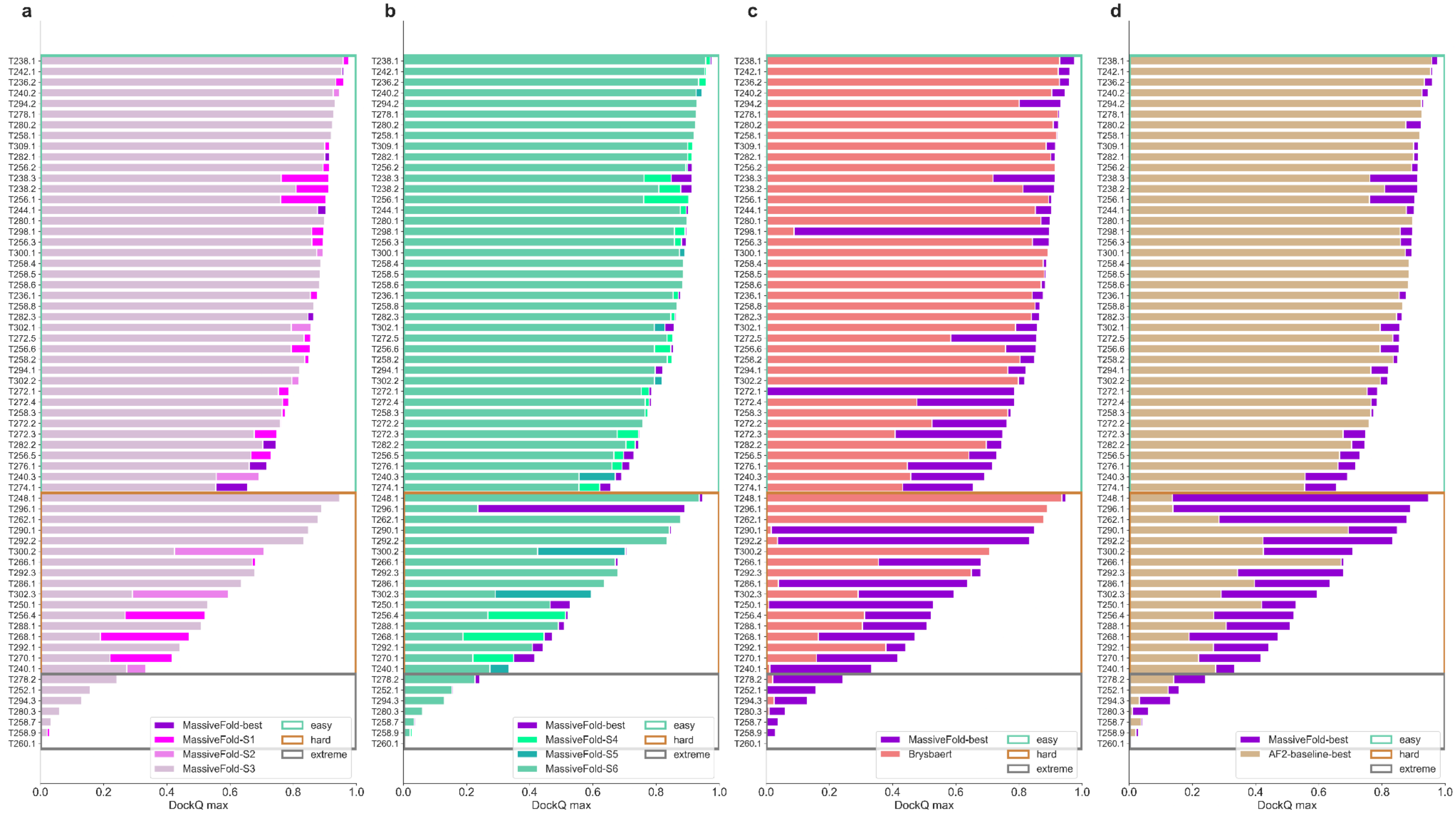


**Figure S5:** Improvement on $DockQ_{max}$ with the best prediction of MassiveFold ('MassiveFold-best') over one or more methods for each interface. These interfaces are gathered by difficulty and further ordered by MassiveFold-best $DockQ_{max}$. **a)** Compares the following massive sampling scenarios used: S1, S2 and S3. **b)** Compares the following massive sampling scenarios used: S4, S5 and S6. **c)** Compares the best prediction in the massive sampling dataset ('MassiveFold-best') with the best among the top 5 structures of the same set ranked by AF2 confidence score ('Brysbaert' submission). **d)** Compares the best prediction in the massive sampling dataset with the best prediction in the basic 25 predictions of AF2v3.
